# NeoPredPipe: High-Throughput Neoantigen Prediction and Recognition Potential Pipeline

**DOI:** 10.1101/409839

**Authors:** Ryan O. Schenck, Eszter Lakatos, Chandler Gatenbee, Trevor A. Graham, Alexander R.A. Anderson

## Abstract

Next generation sequencing has yielded an unparalleled means of quickly determining the molecular make-up of patient tumors. In conjunction with emerging, effective immunotherapeutics for a number of cancers, this rapid data generation necessitates a paired high-throughput means of predicting and assessing neoantigens from tumor variants that may stimulate immune response. Here we offer <u>N</u>eo<u>P</u>red<u>P</u>ipe (Neoantigen Prediction Pipeline) as a contiguous means of predicting putative neoantigens and their corresponding recognition potentials for both single and multi-region tumor samples. NeoPredPipe is able to quickly provide summary information for researchers, and clinicians alike, on neoantigen burdens while providing high-level insights into tumor heterogeneity given somatic mutation calls and, optionally, patient HLA haplotypes. Given an example dataset we show how NeoPredPipe is able to rapidly provide insights into neoantigen heterogeneity, burden, and immune stimulation potential. Through the integration of widely adopted tools for neoantigen discovery NeoPredPipe offers a contiguous means of processing single and multi-region sequence data. NeoPredPipe is user-friendly and adaptable for high-throughput performance. NeoPredPipe is freely available at https://github.com/MathOnco/NeoPredPipe.

## Introduction

Cancer cells are fraught with genomic variants in all regions of the genome with high degrees of heterogeneity in a spatially complex tumor. This intra-tumor heterogeneity (ITH) realizes a fitness landscape upon which natural selection can act (reviewed by [5]). Neoantigens, epitopes derived from proteins translated from non-synonymous variants, are able to make their way to the cell surface in the hopes of stimulating an immune response after a number of cellular processing steps have occurred, primarily proteosomal cleavage and binding with major histocompatibility complexes (MHC) I or II. This binding depends upon the patient specific human leukocyte antigen (HLA) alleles. From here, the bound neoantigen with its MHC-Class I complex makes its way to the cell surface where it may bind with cytotoxic T-cell receptors thereby eliciting infiltration of cytotoxic T-cells capable of detecting and eliminating cells carrying the neoantigen in the absence of immune evading tactics. The immune response is strongly influenced by the total number of neoantigens within a tumor, especially in hyper-mutated cancers ([6]), as well as the ITH of antigenic mutations ([4]). ITH is now being further evaluated using multi-region sequencing approaches whereby adjacent regions of the same tumor or tissue are able to provide greater insights into variant clonality (i.e. truly clonal, subclonal, or shared).

A number of tools have provided means of variant annotation, assessing neoantigen candidacy, and T-Cell receptor (TCR) binding probabilities, but none possess the capability of providing these on multi-region sequence data in bulk or run contiguously as a single tool. Here, we present NeoPredPipe, capable of processing single and multi-region variant call format (VCF) files, carrying out variant annotations, neoantigen predictions, cross-referencing with known epitopes, and performing TCR recognition potential predictions in a single, clear, and proficient pipeline (Figure 1).

**Figure 1.**
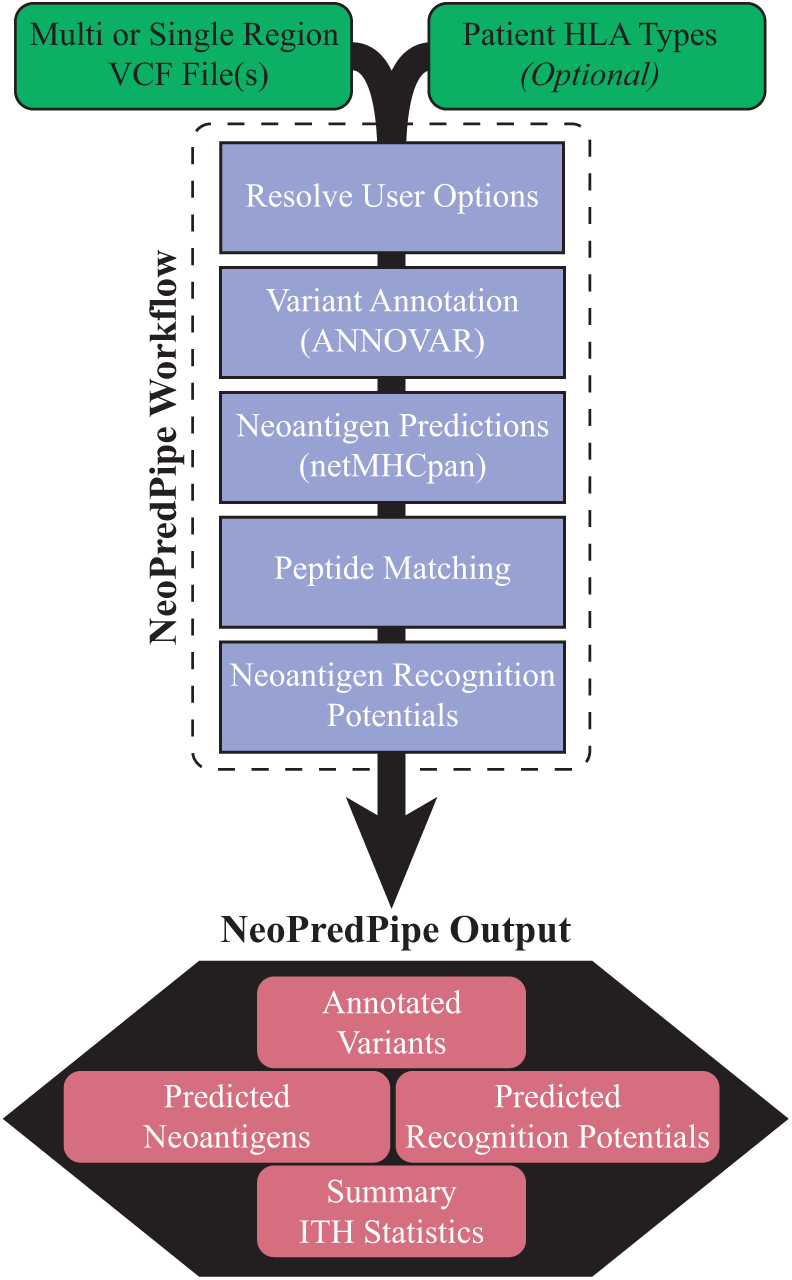
NeoPredPipe workflow differentiating between user steps (green) and execution processes (purple). NeoPredPipe provides low level details and high level summary statistics as output for downstream analysis (red).

## Implementation

The first stage in neoantigen identification from a VCF file is the proper annotation of variants to identify non-synonymous variants. To this end, NeoPredPipe employs the widely used and efficient ANNOVAR ([8]). Specifically, ANNOVAR processes samples in a way that prioritizes exonic variants, this step provides a useful means for quickly partitioning variant calls for downstream applications. The user is able to specify the genome build that they would like to use, provided it is compatible with ANNOVAR. This annotation phase also results in the extracted peptide sequence given the variant base(s) from the annotated variant calls.

Once the VCF files have been annotated and partitioned with ANNOVAR the program determines if HLA haplotypes have been provided by the user containing the HLA-A, -B, and -C haplotypes. NeoPredPipe does not include HLA allele identification as this step in the pipeline is highly dependent upon the source of the data (WES, WGS, targeted gene panels, transcriptome data, or conducted via experimental methods). In cases where no HLA haplotype information is available the most common alleles of each haplotype are assessed; while cases where the HLA haplotypes are homozygous only that HLA haplotype is used for prediction. HLA haplotypes are cross-referenced with available HLA haplotypes prior to executing netMHCpan ([7]) for the primary neoantigen predictions. As with the primary tool, the user is able to specify the epitope to conduct predictions for (typically epitopes of 8-, 9-, or 10-mer lengths). The output from this process yields a single file containing either filtered or unfiltered (dependent on user options) neoantigen predictions with information on the sample possessing the neoantigen and, in the case of multi-region variant calling, a presence/absence indicator for each of the sequenced regions. These predicted neoantigens are then, optionally, cross-referenced with known epitopes utilizing PeptideMatch ([1]), whereby the candidate epitopes are assessed for novelty against a reference proteome that can be supplied by the user as a fasta file (e.g. from Ensembl or UniProt).

The steps outlined above deliver candidate information for neoantigens from provided variant calls that may be presented to cytotoxic T-Cells, however, this does not inform the likelihood of a neoantigen eliciting an immune response (i.e. binds with a TCR). In order to assess the recognition potential we employ the algorithms and process utilized by [3]. The recognition potential is defined as the product of *A* and *R*, where *A* is the amplitude of the ratio of the relative probabilities of binding for the wildtype and mutant epitopes to the MHC-class I molecules, and *R* is the probability that the neoantigen in question will be recognized by a TCR. To define *A* it is necessary to perform neoantigen predictions for the wildtype and mutant epitope, this is not performed by default by NeoPredPipe, but is supplied as an option to employ as a contiguous pipeline. To define *R*, NeoPredPipe utilizes the multistate thermodynamic model employed by [3], which requires alignment scores for each epitope to a curated Immune Epitope Database list of known epitopes (can be refined and updated by the user, but is provided). In order to incorporate the ability to assess ITH in regards to both effective mutations (non-synonymous variants) and neoantigen burdens, NeoPredPipe is capable of handling multi-region VCF files; further these files can be multi-region in only a select number of samples. Thus NeoPredPipe is able to efficiently handle various experimental designs for neoantigen prediction and assessments providing a summary table for downstream statistical and in-depth analysis.

## Results

The output of the pipeline depends largely on the options set by the user, but at the very least, NeoPredPipe provides a single table of putative neoantigens and their predicted binding affinities. With additional options selected it is possible to include, within a single output, whether an epitope matches a reference proteome and the neoantigen’s recognition potential. In additon, for rapid assessment, NeoPredPipe yields summary statistics on the neoantigen burden for each sample as well as information to assess ITH by reporting neoantigen burdens for clonal, subclonal, and shared variants for multi-region samples.

### Use Case

While a small, two sample, multi-region example dataset is provided with the source code for users, we demonstrate the usefulness of NeoPredPipe by applying it to a previously published dataset examining the evolutionary landscape of colorectal tumors [2]. We select two exemplary patient samples (Adenoma 3 and Carcinoma 7 in the original paper) from the dataset, and apply our pipeline using default parameters to evaluate neoantigens in each sample. Figure 2 illustrates the information included in the standard output of NeoPredPipe and potential analysis that can be performed if NeoPredPipe is combined with the output of other standard bioinformatic methods.

**Figure 2.**
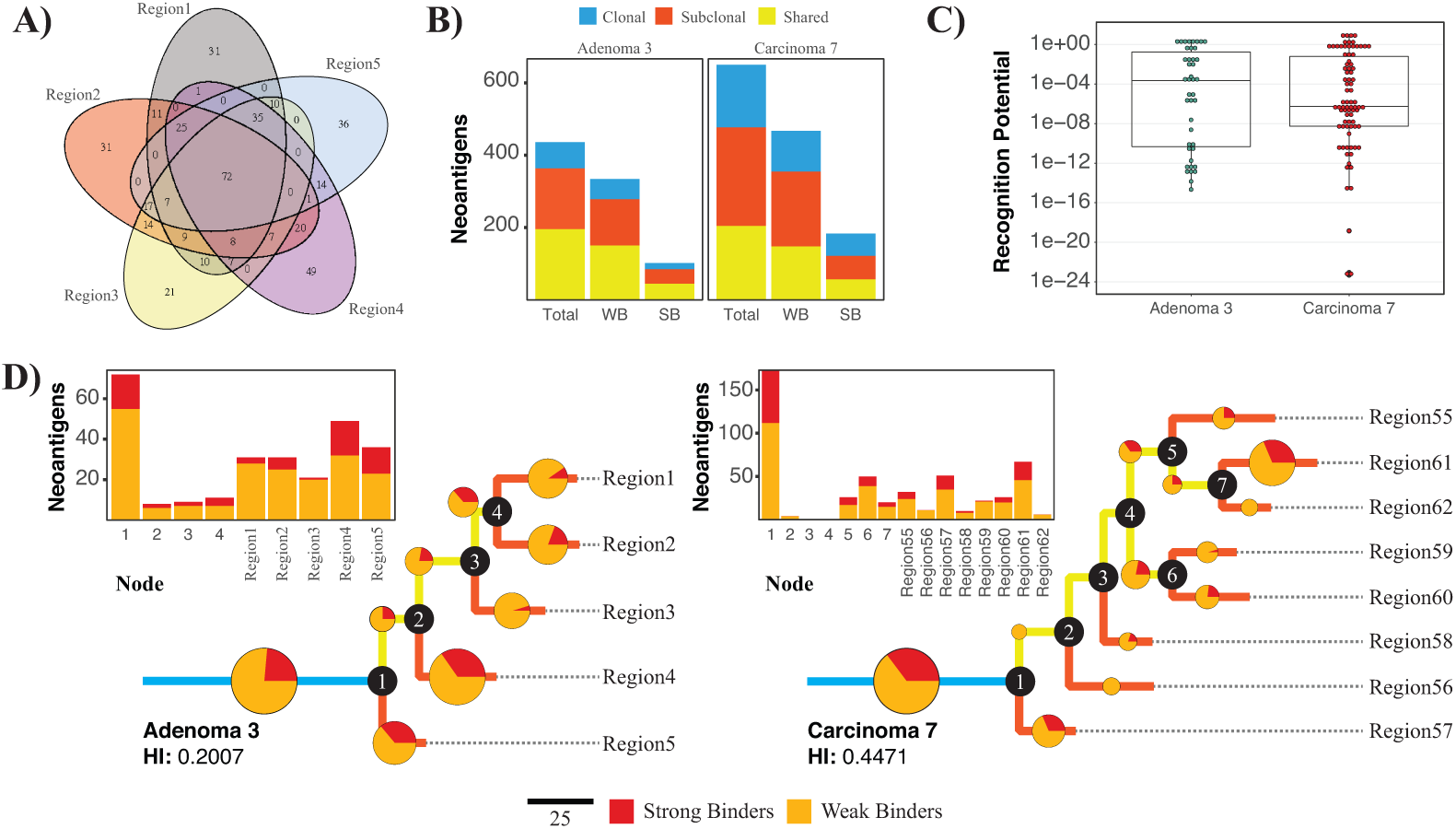
Analysis of neoantigens in two colorectal tumours using NeoPredPipe. (A) Venn diagram of all neoantigens in the five regions of Adenoma 3. (B) Number of neoantigens in the two samples that are clonal (present in all regions, shown in blue), shared (present in at least two regions, in yellow) or subclonal (present in a single region, red). Separate counts of weak and strong MHC-binding neoantigens (WB and SB, respectively) are also shown. (C) Distribution of recognition potential values of neoantigens present in Adenoma 3 (green) and Carcinoma 7 (red). The boxplots represent the median and upper and lower 25 percentile. (D) Phylogenetic tree reconstructed from all exonic mutations for Adenoma 3 (left) and Carcinoma 7 (right). Pie-charts and the bar-charts represent the number of weak (orange) and strong (red) binder neoantigens assigned to each branch. The size of each circle is proportional to the percentage of total of neoantigens on that branch.

Figure 2A provides a summary of the complex interactions between different regions of Adenoma 3, and highlights both Region 4, which harbours the highest amount of subclonal (only present in a single region) neoantigens, and the overall clonality of the sample, with 72 neoantigens detected in all regions. For quick analysis, NeoPredPipe directly outputs a summary of the clonality of neoantigens, also divided into categories of strong and weak binders (peptides with a netMHCpan rank *≤* 0.5 and *≤* 2, respectively). Figure 2B visualizes this summary on two bar-charts for Adenoma 3 and Carcinoma 7. We find that whilst the number of shared neoantigens (present in more than one, but not all regions) is highly similar between the two samples, Carcinoma 7 harbours both more clonal (present in all regions) and subclonal neoantigens; and in total 26% of the neoantigens are clonal, compared to 16% of Adenoma 3. Figure 2C shows the recognition potential value for all neoantigens in the two samples. NeoPredPipe identified 10 peptides in Adenoma 3 and 9 in Carcinoma 7 with a recognition potential value above 1. In Figure 2D, we provide an example of integrating NeoPredPipe outputs with downstream multi-region variant analysis. By inferring phylogenetic trees of each tumor, constructed using all exonic mutations with a variant allele frequency above 0.05 (see [2] for full methods), we find that neoantigen distributions across regions can reflect the phylogenetic distance of regions and clonal structure of samples. 31% and 23.5% of total exonic mutations are clonal in Adenoma 3 and Carcinoma 7, similarly to the clonality of neoantigens shown in Panel B. This approach also highlights regions with neoantigen loads different from their closest neighbors, such as Region61 and Region62 of Carcinoma 7. Therefore the analysis can inform future experimental and bioinformatic investigations of samples allowing for new evolutionary and mechanistic insights into tumor development, evolution, and progression.

## Conclusions

We present NeoPredPipe, an efficient, high-throughput, and user-friendly pipeline for neoantigen prediction and interrogation for single and multi-region tumor VCF files. By tying together commonly utilized bioinformatics toolsets and integrating recent advances in neoantigen assessment, NeoPredPipe yields concise information typically required by researchers and clinicians. Through user options based on computational limitations the pipeline is scalable and customizable for individual research questions. All source code and an extensive read me with all pipeline options are available at https://github.com/MathOnco/NeoPredPipe.

## Availability and requirements

### Project name

NeoPredPipe

### Project home page

https://github.com/MathOnco/NeoPredPipe

### Operating system

Unix-based operating system

### Programming languages

Python and Bash

### Other requirements

Python 2.7, ANNOVAR, netMHCpan, PeptideMatch, and, optionally, NCBI BlastX+.

## Author’s contributions

ROS conceived NeoPredPipe, wrote all scripts, and prepared the manuscript. EL contributed to the writing of the final code base, led debugging efforts, and conceived the use case example. CG provided insights into NeoPredPipe’s necessary outputs. TAG and ARAA provided guidance on code development and oversaw all work efforts. All authors read, edited, and approved the manuscript.

## Acknowledgements

The authors would like to acknowledge William Cross and Ian Tomlinson for sharing their data used in the use case example. ARAA and CG were supported by the U54CA143970 grant from the US National Institutes of Health (NIH) National Cancer Institute (NCI). EL and TAG was supported by Cancer Research UK (grant no. A19771).

